# Integrative Data Analysis to Uncover Transcription Factors Involved in Gene Dysregulation of Nine Autoimmune and Inflammatory Diseases

**DOI:** 10.1101/2024.04.24.591013

**Authors:** Nader Hosseini Naghavi, Steven M. Kerfoot, Parisa Shooshtari

## Abstract

Autoimmune and inflammatory diseases are a group of > 80 complex diseases caused by loss of immune tolerance for self-antigens. The biological mechanisms of autoimmune diseases are largely unknown, preventing the development of effective treatment options. Integrative analysis of genome-wide association studies and chromatin accessibility data has shown that the risk variants of autoimmune diseases are enriched in open chromatin regions of immune cells, supporting their role in gene regulation. However, we still lack a systematic and unbiased identification of transcription factors involved in disease gene regulation. We hypothesized that for some of the disease-relevant transcription factors, their binding to DNA is affected at multiple genomic sites rather than a single site, and these effects are cell-type specific. We developed a statistical approach to assess enrichment of transcription factors in being affected by disease risk variants at multiple genomic sites. We used genetic association data of nine autoimmune diseases and identified 99% credible set SNPs for each trait. We then integrated the credible SNPs and chromatin accessibility data of 376 samples comprising 35 unique cell types, and employed a probabilistic model to identify the SNPs that are likely to change binding probability of certain transcription factors at specific cell types. Finally, for each transcription factor, we used a statistical test to assess whether the credible SNPs show enrichments in terms of changing the binding probability of that transcription factor at multiple sites. Our analysis resulted in identification of significantly enriched transcription factors and their relevant cell types for each trait. The prioritized immune cell types varied across the diseases. Our analysis proved that our predicted regulatory sites are active enhancers or promoters in the relevant cell types. Additionally, our pathway analysis showed that the majority of the significant biological pathways are immune-related. In summary, our study has identified disease-relevant transcription factors and their relevant cell types, and will facilitate discovering specific gene regulatory mechanisms of autoimmune diseases.

## INTRODUCTION

Autoimmune and inflammatory diseases are common complex diseases that affect around 5% of the world’s population [1]. Autoimmunity is initiated when immune self-tolerance fails, resulting in immune attack of different tissues. More than 80 different autoimmune diseases have been reported so far, some of which affect a specific tissue and/or organ, such as Type 1 Diabetes (T1D) which affects pancreas, while others may affect the whole body, such as systemic lupus erythematosus (SLE) that damages a number of organs including joints, skin, brain, lungs, kidneys, and blood vessels [2][3]. The initiating molecular triggers and the underlying immune processes driving each disease can widely differ There is a crucial need to understand the biological mechanisms of autoimmune diseases at cellular and molecular level to both understand the cause of disease and identify therapeutic targets.

Identifying the genetic risk variants that are significantly associated with the diseases is an important step towards the identification of cellular and molecular mechanisms of autoimmune diseases [4]. Genome Wide Association Studies (GWAS) have been successful in identifying hundreds of genomic loci and thousands of genetic risk variants (e.g., single nucleotide polymorphisms or SNPs) associated with increased risk of developing autoimmune diseases [5]. Although most of the disease-associated risk variants have small effect sizes individually, they work together and affect specific biological pathways cumulatively to influence the overall risk of developing a given disease [5][6]. In order to better understand how genetic variants might affect cellular and molecular processes leading to autoimmune diseases, it is essential to develop integrative data analysis methods to integrate multiple data types and to predict the effects of genetic variants on functional elements.

Several studies have reported that the majority of risk variants are found on cell type specific chromatin accessibility sites, and they are likely to alter the expression of specific genes through changing regulatory elements. Since the chromatin must be open and active within a cell for the regulatory site to be available to transcription factor binding, the effects of disease risk variants are usually cell-type-specific. Open chromatin sites can be identified by next-generation sequencing technologies including DNase-I hypersensitivity sequencing (DNase-I-seq) and the assay for transposase accessible chromatin with high-throughput sequencing (ATAC-seq) [5][7][8][9][10][11][12]. Previously, we used bulk open chromatin data in immune-related cell types to identify the specific genetic risk variants and the specific open chromatin sites that are likely to drive risk to nine autoimmune diseases [13]. To extend our studies, in this study we aimed to identify the specific transcription factors that are likely to be relevant to these nine autoimmune diseases.

It is widely acknowledged that transcription factors play critical roles in dysregulation of genes in autoimmune diseases [14–16]. However, we still lack unbiased genome-wide approaches to systematically identify (a) transcription factors whose functions are likely to be affected in autoimmune diseases; (b) genomic locations where bindings of such transcription factors are affected; and (c) the functional mechanisms underlying these effects. Identifying disease-relevant transcription factors facilitates uncovering gene regulatory pathways underlying the disease and will eventually help develop new therapeutic strategies that modulate activities of relevant pathways.

To uncover the transcription factors relevant to autoimmune diseases in an unbiased way, we need to identify the binding profiles of hundreds of transcription factors in different cell types. Chromatin immunoprecipitation followed by sequencing (ChIP-seq) experiments are the standard of the field for measuring binding activities of transcription factors in each cell type. However, it is not feasible to run ChIP-seq on hundreds of transcription factors in several cell types, including different subtypes of immune cells, given the required cost and time. Therefore, the studies which aimed to predict disease-relevant transcription factors based on ChIP-seq data are limited to a small number of transcription factors in a relatively small number of cell types [17,18]. Several computational methods have been developed to predict genome-wide binding of various transcription factors in several cell types using chromatin accessibility data [18,19] [20]. An example is CENTIPEDE [18], which integrates DNase-I-seq and position weight matrix (PWM) data to classify each candidate binding site (e.g., sites that match a certain PWM) according to whether it is bound or not by a specific transcription factor. However, CENTIPEDE and similar methods do not consider disease risk variants and are not designed to identify specific transcription factors related to a disease.

To overcome these challenges, we have adopted a data integration approach, where we first integrate GWAS data of nine autoimmune diseases and the chromatin accessibility data from various cell types to prioritize the genetic risk variants, the relevant cell types, and the candidate risk-mediating open chromatin sites. We then use a probabilistic model integrating DNaseI-seq data with the predefined sequence models to predict if a given genetic risk variant on DNase-I footprints alters the prior odds of binding of specific transcription factors for > 20 folds [17]. These SNPs are called effect-SNPs. Through this model we have been able to verify whether any of the prioritized risk variants is an effect-SNP, meaning that it affects the binding affinity of individual transcription factors; and if so the binding affinity of which transcription factors is likely to be altered by these effect-SNPs. After identifying the relevant transcription factors and the genomic locations where the transcription factor binding can be affected by the risk variants, we use the GREAT pathway analysis to predict the cell-type specific disease-relevant pathway. The advantage of our data integration method is that it takes an unbiased data-driven approach to simultaneously predict (a) the specific disease-relevant transcription factors; (b) the risk variants that may have an effect on the binding probability of these transcription factors; (c) the specific immune cell types in which the transcription factor binding patterns are likely to be affected; (d) the functional activities of the candidate risk-mediating genomic sites; and (e) the regulatory pathways that are likely to be perturbed in these processes.

## RESULTS

### Prioritizing cell types relevant to autoimmune diseases

In our study, we first identified the specific cell types in which the binding profiles of disease-relevant transcription factors are likely to be affected by the autoimmune diseases genetic risk variants. To achieve this, we integrated genetics association data of nine autoimmune diseases with a publicly available dataset by Moyerbrailean et al. that contains a list of matched SNPs and transcription factors and their cell types specificities [17], and employed a statistical test to assess the relevance of each cell type to each autoimmune disease.

The Moyerbrailean et al. dataset contains: (a) a comprehensive list of SNPs from 1000 Genomes Project (https://www.internationalgenome.org/); (b) the transcription factors that their binding affinities are likely to be changed by different alleles of the tested SNPs along with the genomic coordinates of their binding sites; (c) the specific genomic locations of the open chromatin sites where these transcription factors may bind; and (d) the cell types in which each chromatin accessibility site is open. Finally, the SNPs that can significantly change the binding probability of a transcription factor within each cell type were identified and named as effect-SNPs. This dataset was originally generated by the integration of 1000 Genomes Project SNPs with 1372 transcription factors from JASPAR [21] and TRANSFAC [22] databases, and the chromatin accessibility data marked by DNase-I Hypersensitivity Sites (DHS) from 153 tissues, cell types and cell lines generated by the ENCODE Project [23]. We excluded cell lines for our study, and had 374 samples remaining, corresponding to 27 unique cell types.

We obtained the genetics association data for nine autoimmune diseases from the Immunochip database (https://genetics.opentargets.org/immunobase). This included the summary statistics data from the dense genotyping for autoimmune thyroid disease (ATD) [24], celiac disease (CEL) [25], inflammatory bowel disease (IBD) [26], juvenile idiopathic arthritis (JIA) [27], psoriasis (PSO) [28], primary biliary cirrhosis (PBC) [29], multiple sclerosis (MS) [30], rheumatoid arthritis (RA) [31], and Type-I diabetes (T1D) [32]. For each disease, we applied a Bayesian-based fine-mapping approach [13] to identify a list of SNPs that are highly likely to drive risk to disease. These SNPs are called 99% credible interval (CI) SNPs.

Through overlapping the list of 99% CI SNPs for each autoimmune disease with the Moyerbrailean et al. dataset, we measured the proportion of 99% CI SNPs that are likely to change the binding probability of any transcription factor in each cell type (i.e., observed value). We then compared that number to the background expectation, obtained by measuring the proportion of all 1000 Genome SNPs in immunochip data that are likely to change the binding probability of any transcription factor in each cell type (ie., expected value). Fisher’s Exact test was used to assess the enrichment of observed values compared to the background expectation, and a P value of enrichment was assigned to each cell type for each disease (Supp. Table 1).

**Table 1:** The genes involved in the autoimmune disease-related biological pathways and found to be related to autoimmune disease-related transcription factors. This table comprises three columns showing the biological pathways related to autoimmune diseases, their binomial adjusted P values, the genes involved in those pathways, and the disease type.

| Disease | Gene | Pathway (P value) |
| --- | --- | --- |
| IBD | SATB1 | activated T cell proliferation (2.18e-02), CD8-positive, alpha-beta T cell differentiation (4.18e-02) |
| JIA | IRF1 | regulation of MyD88-dependent toll-like receptor signaling pathway (2.41e-12), regulation of alpha-beta T cell proliferation (3.42e-07), positive regulation of interleukin-12 biosynthetic process (3.15e-09), cellular response to interferon-beta (2.22e-09) |
| MS | CLECL1 | B cell adhesion (5.76e-18), interleukin-4 secretion (5.12e-10), positive regulation of mononuclear cell proliferation (5.90e-04) |
| MS | PLEK, CLECL1, ZMIZ1 | positive regulation of cell activation (3.05e-06) |
| MS | ZFP36L2, CLECL1 | leukocyte cell-cell adhesion (1.03e-03) |
| PSO | RUNX3 | peripheral nervous system neuron development (4.61e-06), peripheral nervous system development (1.82e-03), chondrocyte differentiation (3.18e-03) |
| PSO | STX4 | positive regulation of eosinophil degranulation (3.25e-03) |
| PSO | RUNX3, STAT5A, STAT3 | regulation of epithelial cell proliferation (2.92e-02) |
| RA | CD244, CD28, PRKCQ | regulation of alpha-beta T cell activation (4.64e-07), positive regulation of lymphocyte proliferation (1.18e-05), positive regulation of mononuclear cell proliferation (1.19e-05), positive regulation of leukocyte proliferation (1.53e-05) |
| RA | SLAMF7, CD244 | natural killer cell activation (7.36e-09) |
| RA | LY9 | T-helper 17 cell lineage commitment (8.01e-07) |
| T1D | IL7R, SOCS1, PTPN2 | regulation of T cell differentiation (2.67e-03), regulation of T cell activation (1.80e-02), regulation of leukocyte cell-cell adhesion (2.33e-02) |

**Table 2:**
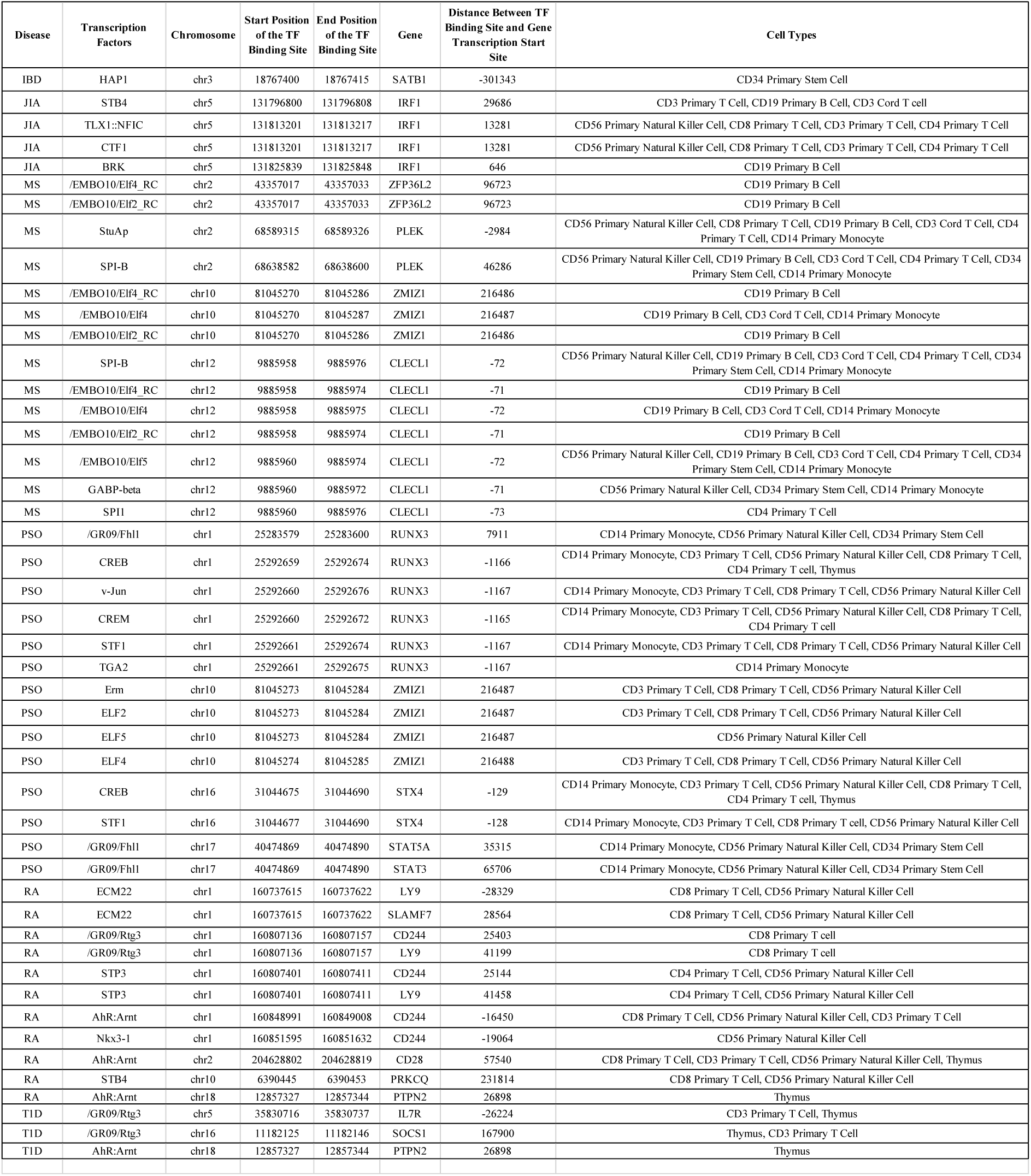
Transcription factor-gene pairs affected by autoimmune diseases risk variants and their cell type activities. This table shows the chromosome, start, end of the binding sites of the affected transcription factors, the cell types comprising those transcription factors, the genes under their control, and their distance to the transcription start site (TSS) of their affected genes.

Except for ATD, CEL and PBC, where no significant cell types were detected, for the other autoimmune diseases, the majority of the relevant cell-types were immune-related (Figure 1, Supp. Figures S1 and S2). The reason we could not find any significant cell types for ATD, CEL and PBC is likely due to their lower GWAS sample sizes; which resulted in a lower number of 99% CI SNPs, and consequently a lower statistical power to detect their relevant cell types. The number of enriched cell types varied between 1 to 7 for the other six autoimmune diseases.

**Figure 1:**
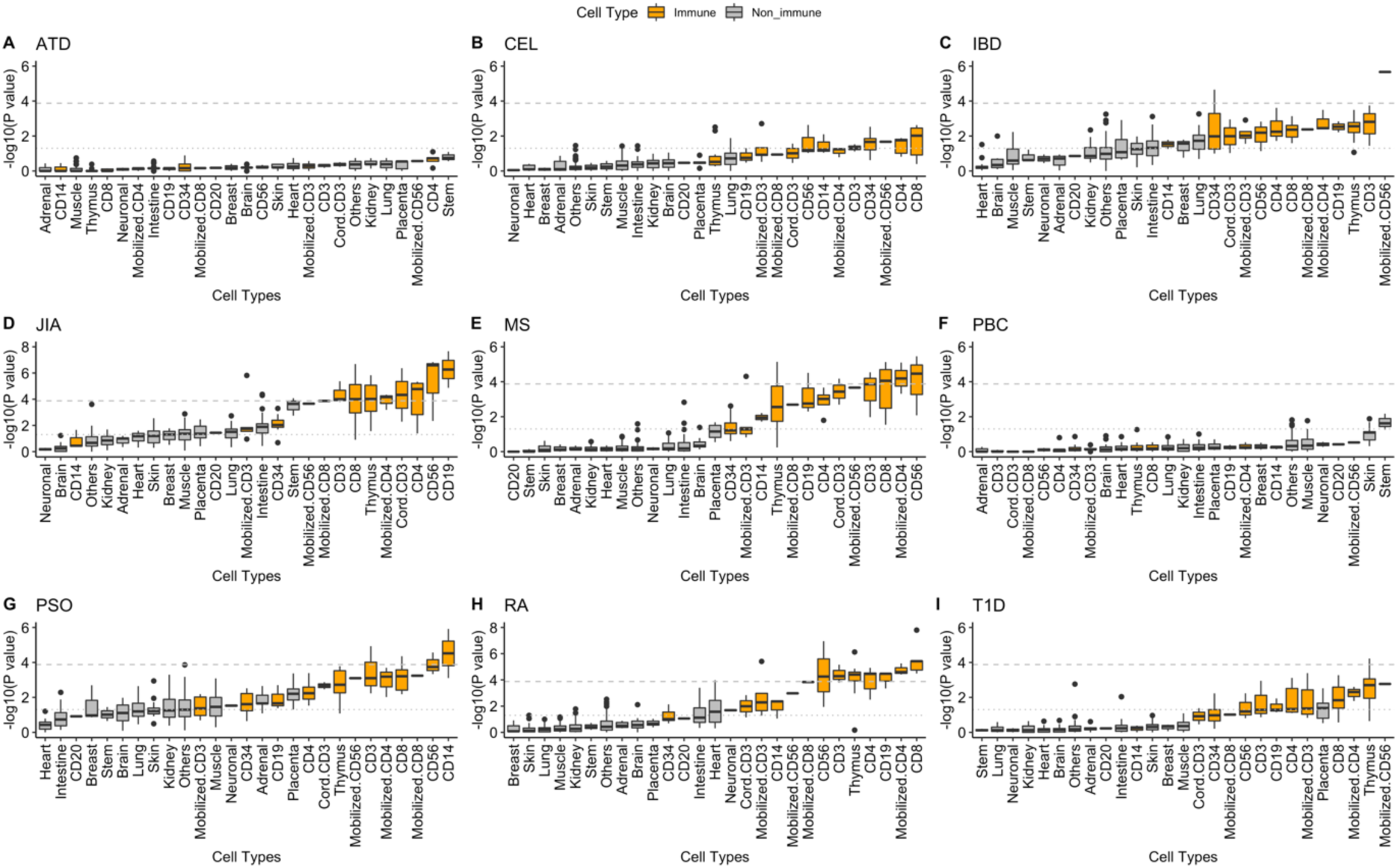
(A-I) boxplots showing the -log10 of original (not corrected) P values of enrichment of samples in nine autoimmune diseases: ATD, CEL, IBD, JIA, MS, PBC, PSO, RA, and T1D. Each box in a boxplot shows -log10 of P values of different samples from a specific cell type. For the majority of the diseases (except ATD, CEL, and PBC) we see that immune cell types are mainly on the right side and some of their samples pass the statistical threshold shown by the dashed lines (-log10 of Bonferroni threshold of 0.05 is shown as the horizontal dashed line, and -log10 of 0.05 is shown as dotted line).

Our analysis identified a sample from common myeloid progenitors (CD34^+^ primary cells) and one sample from CD56^+^ mobilized natural killer cells to be significantly associated with IBD, an observation that is supported by the previous findings that show the relevance of natural killer cells to IBD [33]. Similarly, our analysis identified samples from CD8^+^ alpha-beta T cells, mobilized CD4^+^ helper T cells, CD4^+^ naive T cells, CD3^+^ T cells, CD56^+^ natural killer cells, CD19^+^ B cells and thymus to be highly relevant to RA. This was also supported by previous studies that show the relevance of CD4^+^ T-cells, B cells, and CD56^+^ natural killer cells to RA [34][35]. For MS, we found highly relevant samples from CD56^+^ natural killer cells, mobilized CD4^+^ helper T cells, and CD8^+^, alpha-beta T cells, a finding that is compatible with the previous studies that support the importance of regulatory CD4^+^ T cells, and CD8^+^CD56^-^perforin^+^ T cells in MS [36][37]. For JIA, we found that CD19^+^ B cells, CD56^+^ primary natural killer cells, CD4^+^ naive T cells, CD3^+^ T cells from cord blood, mobilized CD4^+^ helper T cells, thymus, CD8^+^ alpha-beta T cells, and CD3^+^ primary T cells were significantly related. Previous studies have also supported our in-silico findings by showing the relevance of CD19^+^ B cells, CD56^+^ primary natural killer cells, and CD8^+^ T cells to JIA [38][39][40]. Similar to a previous study [41], we discovered that in PSO, samples from CD14^+^ primary monocytes and CD56^+^ primary natural killer cells are significantly enriched. Finally, our enrichment analysis indicated that thymus is significantly associated with T1D. This is also compatible with the previous studies supporting the relevance of thymus to T1D [42]. Overall, our cell type predictions based on the enrichment analysis of SNPs affecting transcription factor bindings in specific cell types align well with previous studies. As the next step we aimed to identify specific autoimmune disease-relevant transcription factors in the immune cell types.

### Identifying autoimmune disease-relevant transcription factors affected in immune cell types

In order to detect the disease-relevant transcription factors in each immune cell types, we first measured the number of overlaps between the CI SNPs and the effect-SNPs of each transcription factor in each sample, and then compared this number to the number of overlaps expected by chance (i.e., background expectation). The background expectation was measured as the number of overlaps between the effect-SNPs and the non-CI SNPs obtained from Immunochip (https://genetics.opentargets.org/immunobase). For each transcription factor, these two measurements were compared against each other using the Fisher’s exact test (Supp. Table S2). This statistical test tells us if the number of observed overlaps between the CI SNPs and the effect SNPs of a transcription factor is significantly higher than what is expected by random chance. For each autoimmune disease, we measured the P values for each transcription factor and each cell type, and the transcription factors with the false discovery rate (FDR) of < 10% were considered to be relevant to the autoimmune diseases in the corresponding samples. Figure 2 shows a heatmap of the enrichment signal (i.e. -log10 P value of enrichment) for the significant transcription factors and their relevant cell types for all the autoimmune diseases. Supp. Figure S3 provides a similar heatmap for each disease individually.

**Figure 2:**
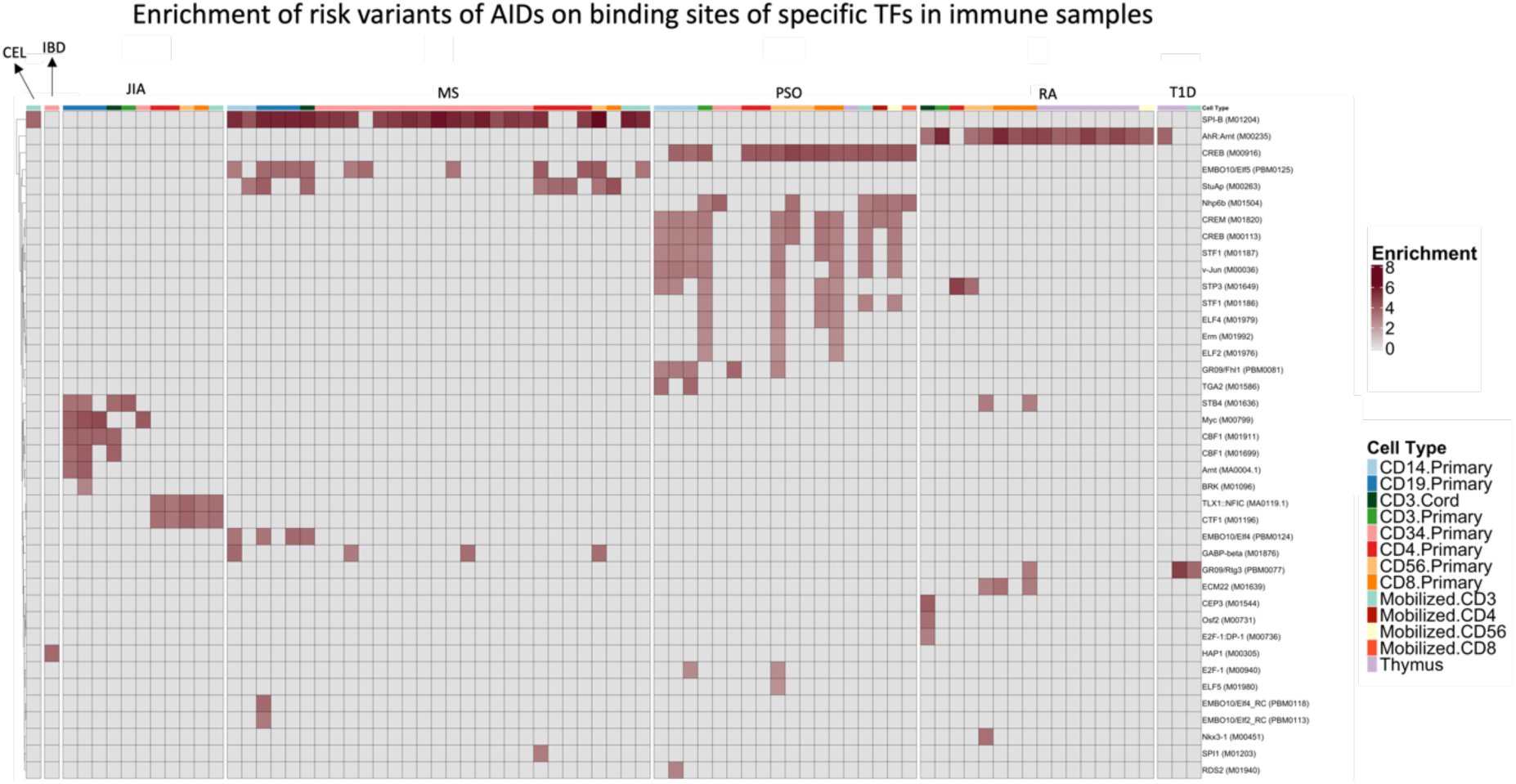
Heatmap of enrichment signal (-log10 of the P value of enrichment) for the transcription factors with FDR < 10% in immune-related samples. Here each column corresponds to one sample, each row corresponds to one transcription factor, and the heatmap colors represent enrichment signal (-log10 of the P value of enrichment). The color bar on the top of the heatmap shows the cell type. The disease name is written on top of the relevant sections of the heatmap. The columns with the same colors from the color bar represent the samples from the same immune cell type. In this heatmap, several transcription factors known to be highly relevant to autoimmune diseases are shown to be significant by our analysis. Examples include Ahr:Arnt for RA, SPI-B for MS, and CREB for PSO.

We calculated the frequency of occurrences of autoimmune disease-relevant transcription factors in immune related samples and cell types (Figure 3). This frequency indicates that each transcription factor is active in how many samples and unique cell types. The number of enriched transcription factors varied between 1 and 16 for seven diseases including RA, MS, PSO, JIA, T1D, IBD and CEL. For two diseases, ATD and PBC, no enriched transcription factor was found (Figures 2 and 3). The GWAS sample sizes for ATD and PBC were smaller than the other seven diseases. This is likely to be the main reason for having a lower statistical power to detect the relevant risk variants, and consequently the relevant transcription factors for these two diseases.

**Figure 3:**
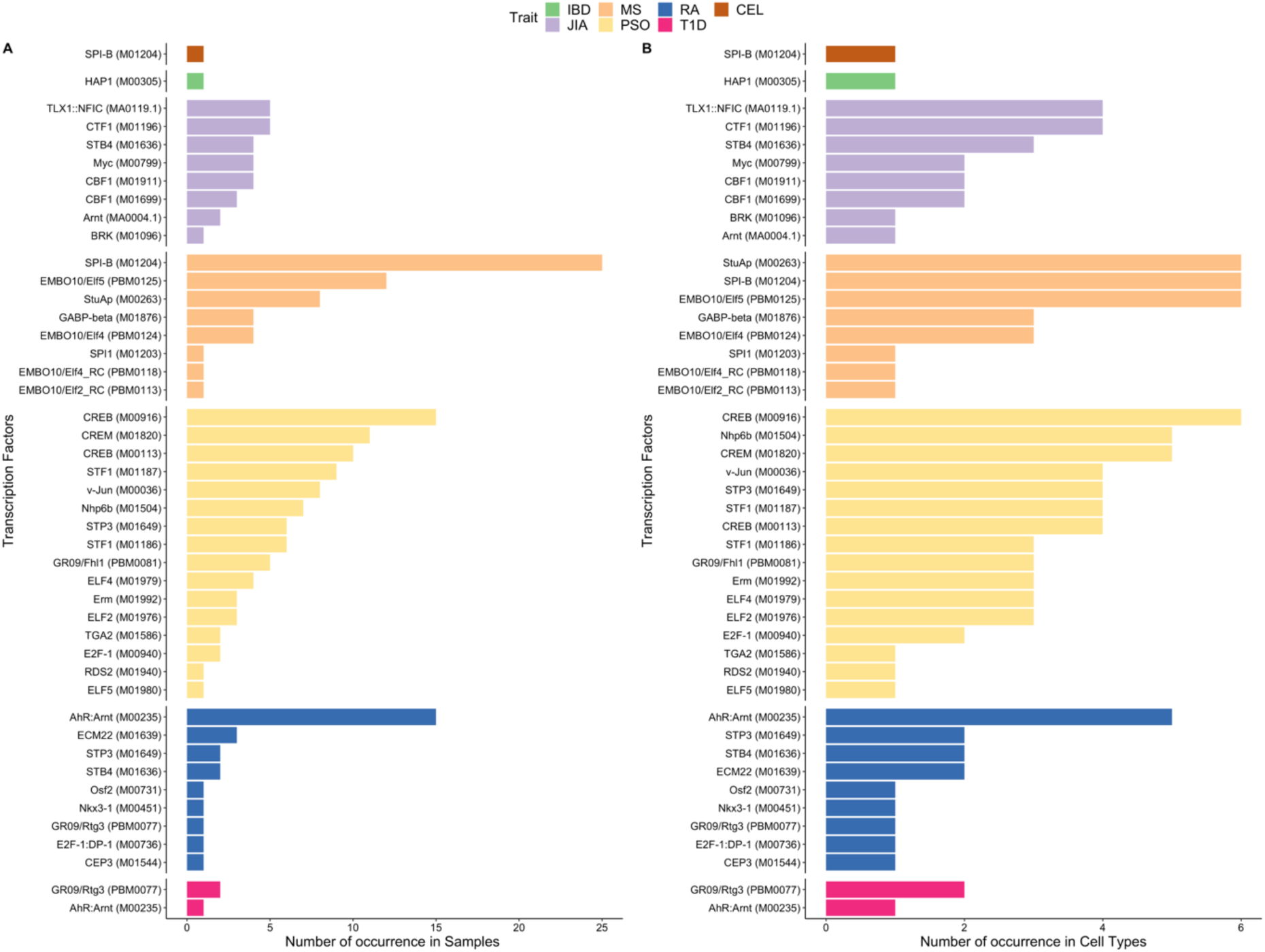
Barplot showing the frequency of occurrence of relevant transcription factors of autoimmune diseases (number of times being active) in samples (A) and unique cell types (B), for each autoimmune disease. Since for each cell type, there exist multiple matching samples, we have shown results for both cell types and samples. Different autoimmune diseases are shown with different colors. For the transcription factors having significant enrichment P values in autoimmune diseases, we also observed high frequency of occurrences in immune samples and cell types. Examples are Ahr:Arnt and SPI-B for RA and MS, respectively.

In RA, we found high enrichment and high frequency values for Ahr:Arnt transcription factor (Figures 2 and 3). The importance of AHR pathway and Ahr transcription factor in RA were discussed in previous studies [43][44]. In our study, a few other RA-relevant transcription factors including ECM22 and STP3 were found in lower frequencies. For MS, we found that the SPI-B transcription factor was highly enriched in a considerable number of immune-related samples (Figures 2 and 3). Recent studies have addressed the relevance of SPI-B to MS [45][46]. The other MS-relevant transcription factors with lower prevalence among immune samples include EMBO10/Elf5, StuAp, and EMBO10/Elf4. For the PSO, we found high enrichment and frequency for CREB, CREM and STF1 transcription factors (Figures 2 and 3). Previous studies have also specified the relevance of CREB and CREM transcription factors to PSO [47][48]. Our study identified other PSO transcription factors with smaller frequencies, including v-Jun, GR09/Fh1, and Nhp6b. For JIA, we detected TLX1::NFIC and CTF1 transcription factors to be highly enriched in multiple immune samples (Figures 2 and 3). This finding is supported by other studies that have shown that CTF1 cytokine (Cardiotrophin 1) is playing a role in JIA [49] [50]. In addition, we found JIA-relevant transcription factors with lower frequencies including Myc, Arnt, CDBF1, and STB4. For T1D, three relevant transcription factors were found in immune samples including GR09/Rtg3 and Ahr:Arnt. For IBD only HAP1 transcription factor was found to be relevant (Figures 2 and 3). Finally, SPI-B transcription factor was found to be relevant to CEL in one immune sample (Figures 2 and 3), while we did not find any sample to be significantly relevant to CEL in the first statistical test (Figure 1).

Overall, this analysis has identified a set of transcription factors relevant to each autoimmune disease. Given that some biological mechanisms could be shared across multiple autoimmune diseases, we asked if some of the prioritized transcription factors are detected in more than one autoimmune disease. We measured the number of common transcription factors and the number of unique cell types with at least one prioritized transcription factor between all pairs of autoimmune diseases (Figure 4). Our analysis showed that two prioritized transcription factors (Ahr::Arnt and GR09/Rtg3) are shared by RA and T1D (Figure 4A). Three transcription factors, STP3, STB4, and SPI-B were shown to be shared between RA and PSO, between RA and JIA, and between MS and CEL, respectively. In terms of the number of commonly relevant STP3, STB4, and SPI-B cell types across multiple autoimmune diseases, we see the highest similarities between JIA, MS, PSO and RA with at least 3 common prioritized cell types between each pair of these autoimmune diseases (Figure 4B).

**Figure 4:**
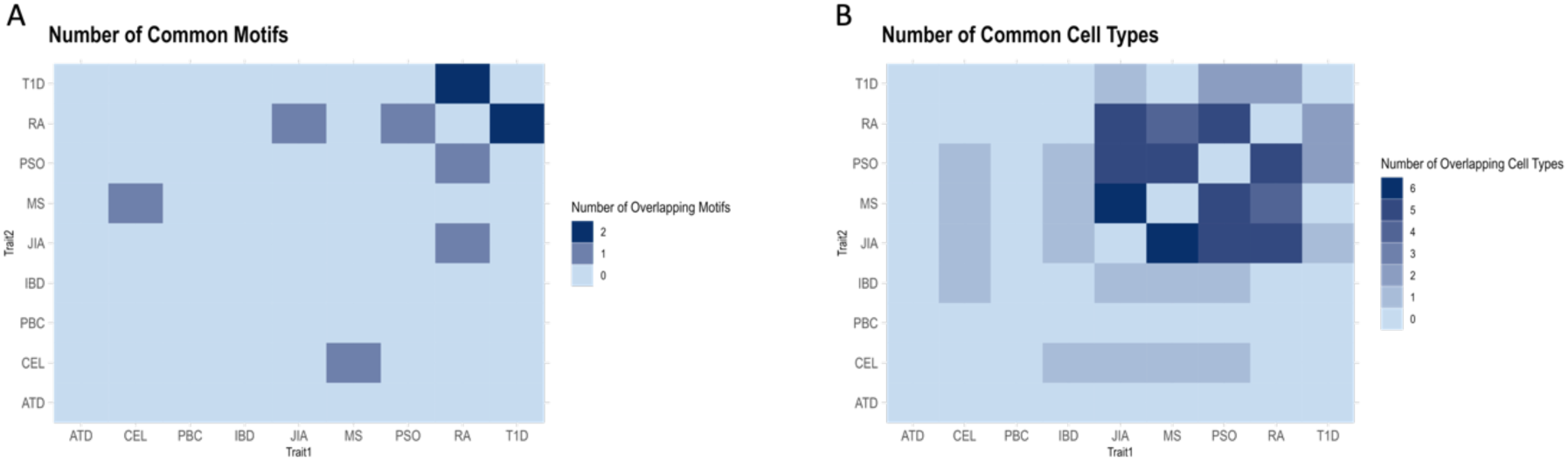
Heatmaps showing the similarities between autoimmune diseases in terms of the number of common transcription factors (A) and the number of common unique cell types (B). In terms of the common transcription factors, there are high similarities between RA and JIA, PSO, T1D and between CEL and MS. In terms of the common unique immune cell types, JIA, MS, PSO, and RA are highly similar.

### Regulatory functions of the prioritized chromatin accessible sites

The open chromatin sites can have different functions, such as enhancer or promoter activity, in a cell-type specific manner. Our analyses identified the disease-relevant transcription factors and the specific open chromatin sites where the binding of the candidate transcription factors is likely to be affected. However, the genomic functions of these candidate disease-relevant chromatin sites were not clear yet. This is mainly because DNase-I hypersensitive sites sequencing is a general assay and it does not provide the specific regulatory function of each site. To address this problem, we used the ChromHMM database [51][52] to identify the regulatory functions of binding sites of the disease-relevant transcription factors in each cell type. ChromHMM uses a reference database to annotate functions of different genomic locations [52]. In order to annotate the genomic functions of the binding sites of candidate transcription factors in our immune-related cell types, we used a set of matched immune cell types from the reference ChromHMM data with the annotated regions from the Roadmap Epigenome project [12] (Supp. Table S3). This enabled the annotation of the genomic functions of the binding sites of autoimmune disease-relevant transcription factors in the relevant immune cell types (Figure 5).

**Figure 5:**
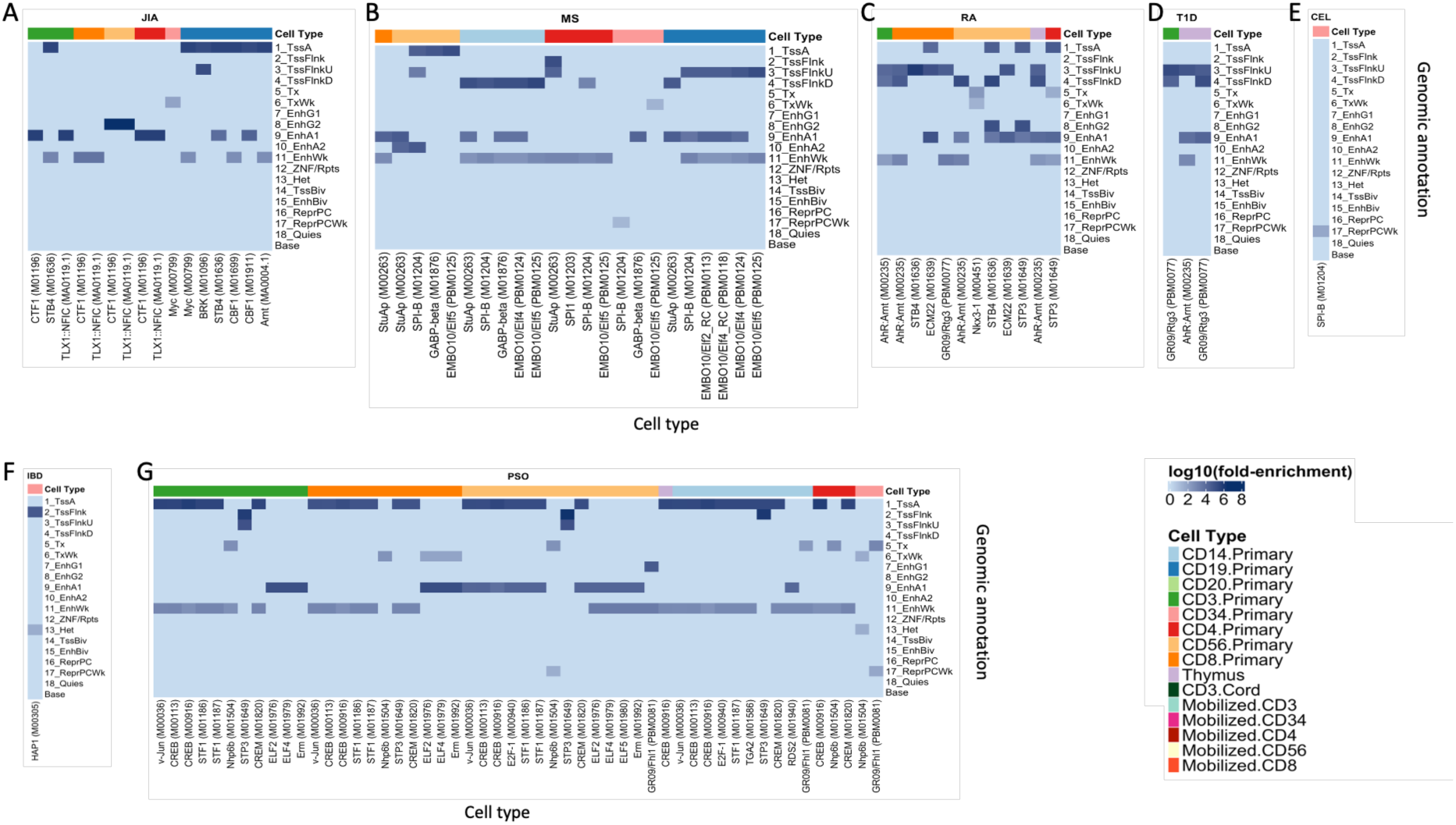
Functional annotations of binding sites of autoimmune disease-relevant transcription factors. The heatmaps show log10 of fold enrichments of each functional annotation on the binding sites of the relevant transcription factors for (A) JIA, (B) MS, (C) RA, (D) T1D, (E) CEL, (F) IBD, and (G) PSO. In these heatmaps, each column represents one transcription factor, and each row corresponds to one functional annotation. The functional annotations include Active TSS (TssA), Flanking TSS (TssFlnk), Flanking TSS Upstream (TssFlnkU), Flanking TSS Downstream (TssFlnkD), Strong transcription (Tx), Weak transcription (TxWk), Genic enhancer 1 (EnhG1), Genic enhancer 2 (EnhG2), Active Enhancer 1 (EnhA1), Active Enhancer 2 (EnhA2), Weak Enhancer (EnhWk), ZNF genes & repeats (ZNF/Rpts), Heterochromatin (Het), Bivalent/Poised TSS (TssBiv), Bivalent Enhancer (EnhBiv), Repressed PolyComb (ReprPC), Weak Repressed PolyComb (ReprPCWk), Quiescent/Low (Quies), and the baseline genomic regions with no specific function (base) [12]. Top colorbar in each heatmap represents the specific immune cell types. Columns with the same color in the top colorbar represent the autoimmune disease-relevant transcription factors being active in the same immune cell type. For the majority of autoimmune diseases (except CEL), binding sites of enriched transcription factors overlap the enhancer regions and/or the transcription start sites and their flanking regions in specific immune samples. This suggests that the prioritized transcription factors primarily bind to the promoters and/or enhancers in an immune cell type-specific manner.

Our analysis indicated that for six out of seven autoimmune diseases with at least one significant transcription factor (i.e., RA, MS, PSO, JIA, T1D and IBD), the binding sites of majority of their relevant transcription factors were significantly located on or near transcription start sites (TSS) and/or enhancer regions of immune cell types. This suggests that the enriched transcription factors mainly bind to promoters and/or enhancers that regulate expression of diseases relevant genes in an immune cell type specific manner. In one autoimmune disease (CEL), however, the enriched transcription factor does not bind to any promoter or enhancer but rather bind to the Weak Repressed PolyComb regions (Figure 5).

Here, we report a few specific cases identified by our analysis. Ahr:Arnt is an RA-relevant transcription factor that we showed to be significant both in terms of enrichment level and frequency (Figures 2 and 3). The binding sites of Ahr:Arnt are significantly overlapping the flanking regions of transcription start sites (TssFlnk*) and enhancer regions (Enh*) in CD3+ primary T cells, CD8+ T cells, CD56+ primary natural killer cells, and thymus. This suggests that Ahr:Arnt binding can be affected at multiple genomic regions including promoters and enhancers that control expressions of RA-relevant genes in specific subpopulations of immune cells (Figure 5, Supp. Figure S4). In the case of MS, we found that SPI-B was relevant both in terms of enrichment and the frequency (Figures 2 and 3). Our integrative analysis using ChromHMM data showed that the binding sites of SPI-B highly overlapped the flanking regions of transcription start sites and enhancer regions in CD56+ primary natural killer cells, CD14^+^ primary monocytes, CD4^+^ naive T cells, and CD19^+^ B cells (Figure 5, Supp. Figure S4). Similarly, for the transcription factors relevant to other autoimmune diseases (except for CEL), we found that their binding sites highly overlap the transcription start sites or their flanking regions, and/or the enhancer regions in specific immune cell types (Figure 5).

All together, these findings support a few findings. (a) The bindings of the autoimmune disease-relevant transcription factors are likely to be perturbed not only at promoters and the transcription start sites, but also at the enhancer sites that regulate genes at a relatively longer distance from them. (b) These effects are cell type specific, and usually multiple immune cell types are relevant. (c) Our analysis can pinpoint specific genomic sites where transcription factors are perturbed, the genomic function of those sites (e.g., enhancer activity), and the cell type specificity of those sites. Supp. Table S4 provides a full list of significant autoimmune diseases risk variants and their genomic locations, affected motifs and transcription factors, relevant samples and cell types, and P values of relevance of motifs for each sample.

### Identifying autoimmune disease-relevant biological pathways

We next aimed to identify the relevant biological pathways for each autoimmune disease. We considered the genomic coordinates of the candidate transcription factor binding sites irrespective of their specific immune cell type activity, and applied GREAT pathway analysis to identify the relevant biological pathways [53]. GREAT pathway analysis considers genes that are close to a set of pre-defined genomic regions, and identifies the relevant biological pathways and their P values of significance [53]. In our case, the pre-defined genomic regions consisted of the candidate binding sites of the relevant transcription factors from all of the immune samples (Supp. Table S4). Figure 6 shows the top significant pathways (FDR < 0.05) for a maximum of 20 pathways per disease. In this figure, the significant pathways are sorted based on their binomial P value of significance.

**Figure 6:**
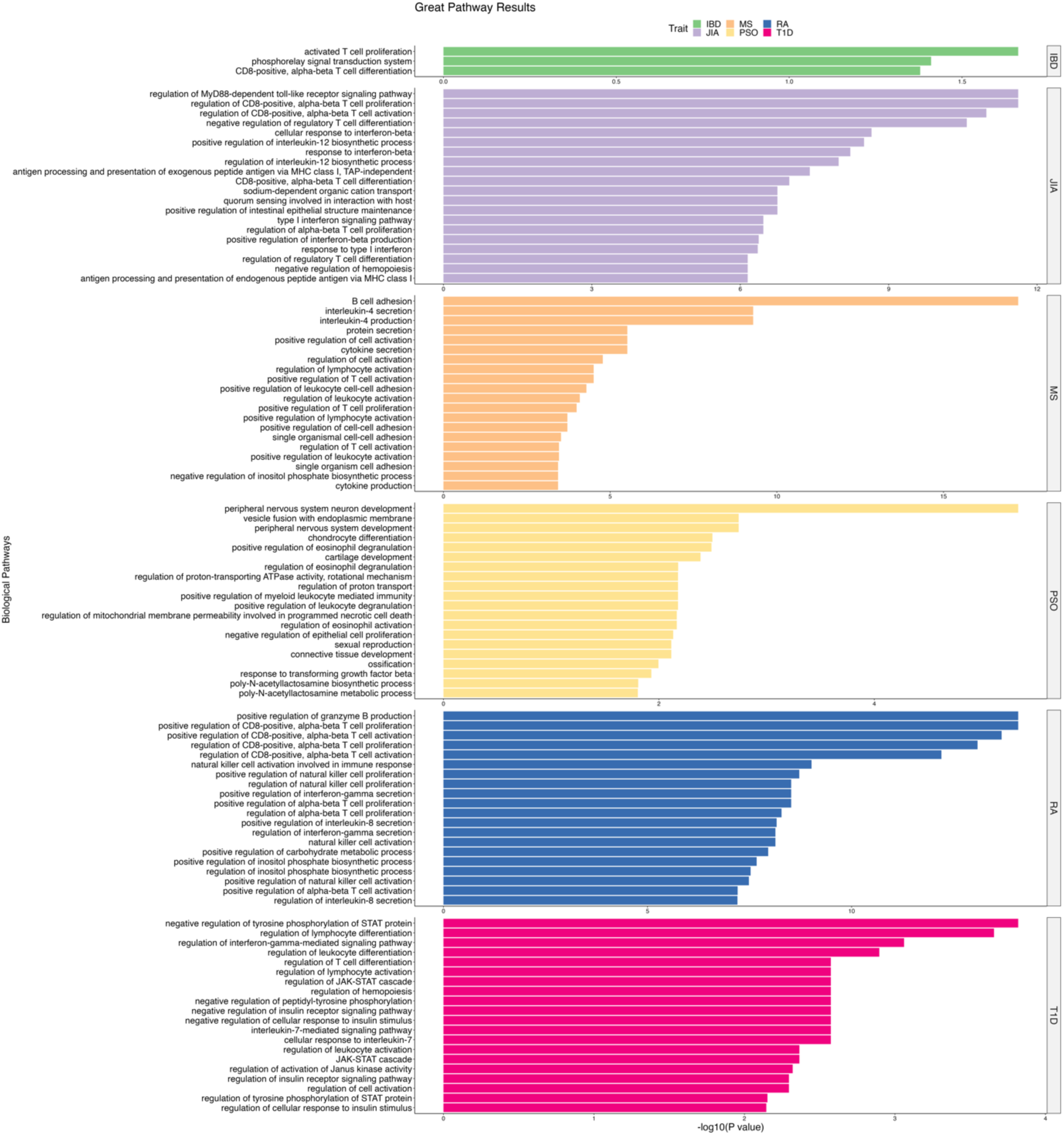
Barplots showing the significant biological pathways (at FDR of 5%) associated with the relevant transcription factors of each autoimmune disease across all immune cell types. Each autoimmune disease is shown with a different color. For the majority of autoimmune diseases with at least one candidate transcription factor (i.e., all diseases except PSO and CEL), the enriched transcription factors in immune cell types are mainly involved in the immune-related biological pathways.

Our GREAT pathway analysis showed that for most autoimmune diseases, majority of the significant pathways are immune-related, with an exception of CEL, ATD, and PBC, for which no significant pathway was detected at the FDR of 1%. More specifically, the significant pathways for RA include positive regulation of CD8^+^ alpha-beta T cell activation/proliferation, regulation of natural killer cell proliferation, and natural killer activation, among several others immune relevant pathways. The relevant pathways for MS included B cell adhesion, interleukin 4 secretion/production, cytokine secretion, and etc. The T1D relevant pathways are related to the regulation of lymphocyte differentiation, interleukin-7-mediated signaling pathway, regulation of leukocyte differentiation, and multiple biological functions involved in the STAT pathway including negative regulation of tyrosine phosphorylation of STAT protein, and regulation of JAK-STAT cascade. For IBD, activated T cell proliferation and CD8^+^ alpha beta T cell differentiation were found to be relevant. For JIA, immune-related pathways including regulation of CD8^+^ alpha beta T cell activation/proliferation and response to interferon-beta were found to be significant (Figure 6). In comparison, for PSO, the majority of pathways were related to blood cell types, skin and nervous system such as regulation of eosinophil degranulation, positive regulation of leukocyte degranulation, peripheral nervous system neuron development, negative regulation of epithelial cell proliferation, and connective tissue development (Figure 6). PSO is an autoimmune disease affecting the skin, and therefore, finding skin-relevant pathways is not unexpected for this disease. In addition, researchers have uncovered the role of the peripheral nervous system in PSO [54].

### Detecting the disease-relevant genes in autoimmune disease-relevant pathways

To further investigate the effect of transcription factors in autoimmune disease-relevant biological pathways, we used utilities provided in the GREAT toolbox, and extracted the genes that GREAT linked to the candidate binding sites of autoimmune disease-relevant transcription factors [53]. This way we identified the affected genes and the transcription factors that bind to the regulatory sites linked to these genes. Supp. Table S5 lists the significant pathways (FDR < 1%), their corresponding FDR, the genes linked to candidate transcription factor binding sites for each of the pathways, and the name of each autoimmune disease. Table 1 highlights some of these cases for a quick review.

To extend our discoveries, we identified the transcription factors linked to each of the genes and the distance between the binding sites of those transcription factors and the transcription start sites of the genes linked to them. If the distance is < 2,000 bp, the transcription factor is considered to be affecting the promoter of the linked gene. Otherwise, the transcription factor is likely to dysregulate the target gene through another gene regulatory activity such as enhancer activity. Supp. Table S6 shows the gene-transcription factor pairs affected by autoimmune diseases risk variants and their cell type activities. This table includes the genomic coordinates (chromosome, start, and end position) of binding sites of the affected transcription factors, names of the transcription factors and their linked genes, and the genomic distance between them. Table 2 highlights some of these cases.

As shown in Supp. Table S5 and Table 1, CLECL1 and PLEK genes are highly involved in the majority of the top MS-related pathways. CLECL1 is highly expressed in dendritic cells [55] and B cells [56], and is known to be involved in the co-stimulation of T cells [57]. Previous studies have confirmed the role of CLECL1 [58] and its role in co-stimulation of T cells by B cells in MS [57]. Our results show that this gene is likely to be regulated by the SPI-B transcription factor in multiple cell types including CD56^+^ Natural Killer cells, CD19^+^ B cells, CD4^+^ T cells, and CD14^+^ Monocytes. More specifically, SPI-B binds to a region close to the transcription start site of the CLECL1 gene (distance of 73 bps) supporting that SPI-B binds to a promoter region of this gene (Supp. Table S6, Table 2).

Our analysis indicates that CD244 is likely to be relevant to RA, and multiple similarly-related pathways are enriched, including regulation of alpha-beta T cell activation/proliferation (Table 1 and Supp. Table S5). We show that three transcription factors, Ahr:Arnt, STP3 and Nkx3-1, are likely to regulate CD244 through enhancer activities (distance > 5,000 bp). These patterns are observed in CD3^+^, CD4^+^, and CD8^+^ T cells and CD56^+^ Natural Killer cells (Tables 2 and Supp. Table S6). CD244 encodes a protein found on the surface of multiple immune cell types including T cells, Natural Killer cells, and Monocytes. CD244 protein regulates the immune response by sending regulatory signals to the nearby cells [59].

Our analysis predicted that PTPN2 and IL7R are involved in the top T1D-related pathways, including regulation of T cell differentiation, regulation of leukocyte cell-cell adhesion, etc. (Tables 1 and Supp. Table S5). PTPN2 encodes a protein playing a significant role in regulation of Interferon Signaling and Endoplasmic Reticulum Stress Response pathways [60]. Dysregulation of these pathways disrupts the function of B cells in the pancreas which leads to T1D [60]. IL7R is also an immune-related gene which encodes the receptor for IL-7 cytokine. IL-7 cytokine is known to have a significant effect on the development of T and B cells through affecting JAK-related pathways [61].

For JIA, IRF1 gene was found repeatedly in multiple relevant pathways, including regulation of MyD88-dependent toll-like receptor signaling pathway, regulation of alpha-beta T cell proliferation/activation (Tables 1 and Supp. Table S5). Previous studies have shown that this gene has a long-range connection to the regions containing JIA risk variants supporting that IRF1 is likely to be affected by the JIA risk variants through enhancer activity [62]. Our study supports the same process.

Finally, our analysis indicated that RUNX3 and STAB1 are involved in the top related pathways of PSO and IBD, respectively (Tables 1 and Supp. Table S5). RUNX3 gene plays a role in immune cell differentiation and activation through affecting myeloid cell lineage [63]. STAB1 is expressed in regulatory T cells and help in immune tolerance. Dysregulation of this gene can lead to autoimmune disorders. STAB1 gene is also expressed in hematopoietic stem cells and plays a role in lymphocyte lineage [64].

## DISCUSSION

While it is widely acknowledged that transcription factors play critical roles in dysregulation of genes in autoimmune diseases [65,66][65,67][65,68][65,69], we still lack unbiased genome-wide approaches to systematically identify (a) the transcription factors whose functions are likely to be affected in autoimmune diseases; (b) genomic locations, where bindings of such transcription factors are affected; and (c) the functional mechanisms underlying these effects. In this study, we have addressed this knowledge gap through in-silico identification of the autoimmune disease-relevant transcription factors. This facilitates uncovering gene regulatory pathways underlying the disease and will eventually help develop new therapeutic strategies that modulate activities of disease-relevant pathways.

Our study is based on the observation that autoimmune diseases risk variants are enriched on open chromatin sites of immune cell types. This supports that disease risk variants are likely to change binding patterns of the transcription factors that bind to the specific chromatin accessible sites, and as a result dysregulate the target genes controlled by these sites in a cell type specific manner. Here, we have taken a statistical-based data integration approach, where we integrate genetics association data of nine autoimmune diseases; with the chromatin accessibility data of 35 cell types and tissues [70] and sequence-based transcription factors databases [21] [22]. For each disease and each cell type, we identified those transcription factors that their binding to DNA is likely to be affected by multiple disease risk variants at multiple genomic locations. We also identified the relevant immune cell types, the regulatory function of the disease-relevant transcription factors, and the affected biological pathways.

Our cell type prioritization analysis confirmed the relevance of multiple immune cell types including CD4^+^ and CD8^+^ T cells in RA, CD19^+^ B cells in MS, CD19^+^ B cells in JIA, as suggested previously. In addition, we found some other cell types that were less studied for the autoimmune diseases of interest, including CD19^+^ B cells and Thymus for RA, CD56^+^ natural killers for MS, CD3^+^ T cells from cord blood and Thymus cells for JIA. One advantage of our method compared to other similar studies [20,71–75] is that our method can directly link the transcription factor binding profiles in 374 samples from 35 unique cell types to the autoimmune diseases risk variants. We have detected the transcription factors that their binding sites overlap with the disease risk variants, and their binding patterns are affected by the risk variants in specific immune cell types. This has enabled us to identify autoimmune disease-relevant transcription factors and their cell type activities simultaneously without requiring to profile activities of hundreds of transcription factors in several cell types using assays such as ChIP-seq.

Several transcription factors prioritized by our analysis were compatible with findings of other studies. This includes Ahr::Arnt for RA and SPI-B for MS. We also identified several novel transcription factors for autoimmune diseases, including StuAp, Elf, and EMBO10 for MS, and ECM22 and STP3 for RA. Our ChromHMM analysis showed that the transcription factors predicted to be relevant to autoimmune diseases mainly bind to the promoter and enhancers of the genes involved in the immune-related pathways. An example is a MS risk variant that affects binding probability of SPI-B transcription factor to the promoter region of *CLECL1* gene. Our GREAT pathway analysis showed that this gene is involved in various enriched immune-related biological pathways including B cell adhesion, interleukin 4 secretion and production, and positive regulation of T cell activation. Similar patterns were found for other autoimmune disease-relevant transcription factors.

In our study, we used bulk DNase-I-seq data of tissues and cell types from healthy individuals and integrated them with genetics association data from patients to make predictions regarding disease-relevant transcription factors and gene regulatory pathways [70]. Chromatin accessibility data from patients are scarce. As the chromatin accessibility data from sequencing technologies (e.g., DNase-I-seq and ATAC-seq) obtained from both healthy individuals and patients with autoimmune diseases become available, a similar approach can be taken to experimentally validate the role of specific transcription factors in the disease. Furthermore, as chromatin accessibility data from single-cell sequencing technology become more widely available, bulk sequencing data can be replaced by the single-cell chromatin accessibility data. This will capture the high heterogeneity among immune cells and will help identifying sub-populations of major immune cell types relevant to autoimmune diseases and their gene regulatory programs.

## METHODS

In our study, we aimed to use publicly available datasets to expand our understanding of cellular and molecular mechanisms of nine autoimmune diseases, with a primary focus on the cell-type-specific effects of disease risk variants on transcription factor bindings [9][10][76][70]. This required the development of computational methods to predict both disease-relevant cell types, and the disease-relevant transcription factors that their bindings are likely to be affected in specific immune cell types [18,70]. Here, we integrated three data types: (a) genetics association data; (b) chromatin accessibility data generated by DNase-I sequencing; and (c) sequence-based transcription factor databases. We applied two different enrichment analyses on these datasets to prioritize disease-relevant cell types and the relevant transcription factors. We also identified the regulatory functions of the binding sites of the relevant transcription factors, and biological pathways relevant to each disease.

### Cell-type-specific transcription factor data

To uncover autoimmune disease-relevant transcription factors and the cell types in which these transcription factors may be affected, one needs to know the binding profiles of hundreds of transcription factors across various cell types. ChIP-seq experiments are the standard of the field for measuring binding activities of transcription factors. However, it is not feasible to run ChIP-seq on hundreds of transcription factors across hundreds of cell types. In addition, identifying the transcription factor activities alone is not sufficient to assess the effects of disease risk variants on transcription factor binding. To address this, we have used a publicly available dataset generated by Moyerbrailean et al. and integrated them with the genetics association data from nine autoimmune diseases to predict disease-relevant transcription factors and their cell-type-specific activities. This large-scale dataset contains the binding sites of 1,372 transcription factors motifs across 153 tissues and cell types, and a list of effect-SNPs. Effect-SNPs are a set of SNPs which are likely to affect the prior odds of binding of specific transcription factors in these sites for > 20 folds as predicted by Moyerbrailean et al. model [17]. Here, we first removed all of the cancer cell lines from the Moyerbrailean et al. dataset since we were mainly interested in analyzing the effects of disease risk variants on transcription factors bindings in primary cell types and tissues. This resulted in a set of 374 samples from 27 unique cell types, including samples from different immune cell types, stem cells, heart, lungs, brain, muscles, brain, placenta, thymus, intestine, breast and adrenal gland.

### Genetics association data

We used Immunochip data (https://genetics.opentargets.org/immunobase) to obtain genetics association data of nine autoimmune diseases including autoimmune thyroid disease (ATD) [24], celiac disease (CEL) [25], inflammatory bowel disease (IBD) [26], juvenile idiopathic arthritis (JIA) [27], psoriasis (PSO) [28], primary biliary cirrhosis (PBC) [29], multiple sclerosis (MS) [30], rheumatoid arthritis (RA) [31], and Type-I diabetes (T1D) [32] (See Supp. Table S7). Immunochip is an Illumina Infinium genotyping chip which captures > 195,000 SNPs, primarily located on the genomic loci with significant association to autoimmune diseases. The immunochip is commonly used in the studies that aim to identify the genetic risk variants of autoimmune diseases and their downstream effects [77][78][79][80]. The main advantages of using Immunochip data for studying multiple autoimmune diseases include: (1) the dense genotyping of autoimmune disease-associated loci is available. This increases the likelihood of identifying disease risk variants, without worrying about SNP imputation; (2) the same chip is used for multiple autoimmune diseases. This facilitates comparing the results across multiple autoimmune diseases in a consistent way without worrying about systematic biases; and (3) the summary statistics data are publicly available for these diseases.

While some of the disease-associated loci can be shared across different autoimmune diseases, there are still variabilities across diseases both in-terms of the associated genomic loci and the disease risk SNPs in those loci. To ensure we are targeting the right associated loci for each disease, we considered the largest GWAS dataset for each disease and compiled their list of lead SNPs (i.e., most associated SNPs per locus) [24][25][27][30][28][81][32][82][83]. We then overlapped the lead SNPs from the largest GWAS datasets with the SNPs from the immunochip data. For each disease, we only kept those immunochip loci that had an overlapping lead SNP from the largest GWAS dataset for the disease. We then applied a Bayesian fine-mapping approach to identify a set of 99% credible interval (CI) SNPs for each risk locus [7,84]. The CI SNPs of each association locus are a set of SNPs that together are likely to include the causal SNPs within them with a 99% chance, if the causal SNP is genotyped in the immunochip data.

### Enrichment analysis for cell type identification

We prioritized cell types in which the transcription factor bindings are likely to be affected in each autoimmune disease, through assessing the enrichment of disease risk variants on the transcription factor binding sites of various cell types and tissues. For our enrichment analysis, we grouped the SNPs into two sets, named as test and null SNPs. The test SNPs include the 99% CI SNPs that were obtained by using the Bayesian-based fine-mapping approach for each autoimmune disease [84], and the null SNPs are defined as the set of Immunochip SNPs that are not in the list of CI SNPs of the disease. For each of the 374 samples from 27 unique cell types, we measured the number of SNPs falling into the test (i.e., CI SNPs) and null (non-CI SNPs) groups, and asked how many of them tend to be an effect-SNP in that sample. Here, Effect-SNPs are a set of SNPs that are predicted to change the binding probability of any transcription factor > 20 folds based on Moyerbrailean et al. model [17] (Supp. Table S1). For each disease and each sample, we compared the proportion of effect-SNPs within the test and null SNPs, and obtained a P value of enrichment using Fisher’s Exact test. We adjusted the P values for multiple testing using Bonferroni correction, and identified samples with adjusted P values less than 0.05. A significant adjusted P value is an indication that the disease CI SNPs are affecting transcription factor bindings in a sample more likely than expected by random chance. This results in prioritization of samples with significant P values as likely candidates for the disease-relevant cell types. Since for each cell type and tissues, we usually had multiple samples, we generated boxplots of -log10 (p values) and showed the reproducibility of the results across samples from the same cell types (Figure 1).

### Enrichment analysis for identifying autoimmune disease-relevant transcription factors

Next in our analysis, we aimed to identify the specific transcription factors that are relevant to each autoimmune disease in the disease-relevant cell types. Since our enrichment analysis for cell type identifications prioritized immune cells as autoimmune disease-relevant cell types, we focused the rest of our analysis on the immune cell types. Here again we used Fisher’s Exact test, but this time, instead of using all the effect-SNPs of a sample, we considered the effect-SNPs relevant to each individual transcription factor separately. In the Moyerbrailean et al. dataset [70], some transcription factors have multiple motifs. We pooled all motifs of each transcription factor. We named a transcription factor likely to be "active" in a sample, if it is bound to at least one binding site in that sample according to the Moyerbrailean dataset. Similar to before, we grouped SNPs into test (i.e., CI SNPs) and null (i.e., non-CI SNPs) groups. Then for each immune-related sample, we extracted a list of effect-SNPs of each "active" transcription factor in that sample, and measured how many test and null SNPs are also an effect-SNPs (Supp. Table S2). Then for each sample-transcription factor pair, we applied the Fisher’s Exact test to assess the enrichment of disease CI SNPs in terms of being an effect-SNP for a specific transcription factor in a specific cell type. We used the False Discovery Rate (FDR) threshold of 0.05 to identify the significant transcription factors for each immune-related sample. This resulted in the prioritization of the transcription factors that their binding sites are likely to be affected in each immune cell type.

### Discovering the gene regulatory functions of the affected transcription factors

Transcription factors play a vital role in regulation of gene expressions in cells. One mechanism through which the disease genetic risk variants are likely to affect regulations of specific genes is through changing the binding patterns of specific transcription factors in certain cell types. In our study, we used Fisher’s Exact test to identify transcription factors that their bindings are likely to be affected by disease-associated risk variants in multiple chromatin accessible sites open in immune cell types. We overlapped SNPs positions with the transcription factors binding sites to identify the specific genomic regions, where the bindings of the relevant transcription factors are likely to be affected by the disease risk variants. As the next step of the analysis, we aimed to identify the genomic functions of the risk-mediating regulatory sites, which are the sites where binding of the affected transcription factors is likely to be affected. DNase-I sequencing is a general mark for regions of open chromatin, where transcription factors can bind to DNA. However, DNase-I-seq does not identify the genomic function and activity (e.g., enhancer or promoter activity) of these regulatory sites. To address this, we used ChromHMM toolbox [51][52] and annotated the regulatory functions of the risk-mediating regulatory sites. ChromHMM annotates specific genomic regions of various cell types based on a reference database [51][52]. We obtained annotations from reference immune-related cell types using the Roadmap Epigenomics project [12]. For each cell type, we considered the matched cell types from this reference database. ChromHMM measures the fold enrichment of each annotation type (e.g., enhancer, promoter, transcription, etc.) from the reference cell types on the genomic regions ((here, binding sites of the affected transcription factors) of the cell types of interest [85], and generates an annotation by cell type matrix with significant fold enrichment values. The enrichment values < 2 are set to zero [51][52]. We first extracted the binding sites of disease-relevant transcription factors in immune-related cell types (Supp. Table S4). Then we used the functional annotations of the reference cell types from the Roadmap Epigenome dataset [12], and matched the cell types from the reference database with the immune cell types in our datasets based on the gene marker lists (Supp. Table S3). We then used ChromHMM to annotate the binding sites of active transcription factors in different cell types.

### Applying pathway analysis on enriched transcription factors of each autoimmune disease

We used the GREAT pathway analysis tool to find the biological pathways, as related to the disease-relevant transcription factors. We first extracted the genomic regions, where the binding of disease-relevant transcription factors can be affected by disease risk variants. While some of these sites overlap gene promoters, some other sites are located on the intergenic regions, and may have enhancer activity. The GREAT pathway analysis tool links the genomic regions (here, binding sites of the affected transcription factors) with the genes, and then performs pathway analysis on the linked genes [53]. In addition to identifying the relevant pathways, GREAT identifies a set of genes that were linked to each pathway.

### Identifying the shared transcription factors between pairs of autoimmune diseases

We assessed the similarities between pairs of autoimmune diseases, in terms of the shared transcription factors and cell types in two ways. For each pair of autoimmune diseases, we first measured the number of common prioritized transcription factors across all immune samples. We then identified those cell types that have at least one disease-relevant transcription factor, and measured the number of common cell types for each pair of autoimmune diseases.

## DATA AVAILABILITY

DHS data of all samples along with the motif data and effect SNPs information can be downloaded from http://genome.grid.wayne.edu/centisnps/#files. The reference annotation files for the ChromHMM analysis can be downloaded from the Roadmap Epigenome Project website: https://www.ncbi.nlm.nih.gov/geo/roadmap/epigenomics/. ChromHMM package is available at http://compbio.mit.edu/ChromHMM/, and GREAT pathway analysis can be obtained from http://great.stanford.edu/public/html/. The Immunochip data of autoimmune diseases can be obtained from the Immunobase portal https://genetics.opentargets.org/immunobase or from the reference papers (see Supp. Table S7). All of the codes used in our analysis are available at the GitHub link: https://github.com/shooshtarilab/AID-TF.

## ACKNOWLEDGMENTS

The authors would like to acknowledge Digital Research Alliance of Canada for providing high-performance computing resources for this project. We thank Dr. Aidin Foroutan Naddafi for editing the first draft of the paper. This project was supported by the Government of Canada through the Natural Sciences and Engineering Research Council (NSERC) (RGPIN/04062-2021 and DGECR-2021-00298) and the internal research funds from Western University, Schulich School of Medicine and Dentistry, Children’s Health Research Institute, and Lawson Health Research Institute. PS is also supported by an Early Investigator Award from the Ontario Institute for Cancer Research (OICR). The funders had no role in the study design, data collection and analysis, decision to publish, or the preparation of the manuscript.

