## Supplementary Figures for "Integrative Data Analysis to Uncover Transcription Factors Involved in Gene Dysregulation of Nine Autoimmune and Inflammatory Diseases"

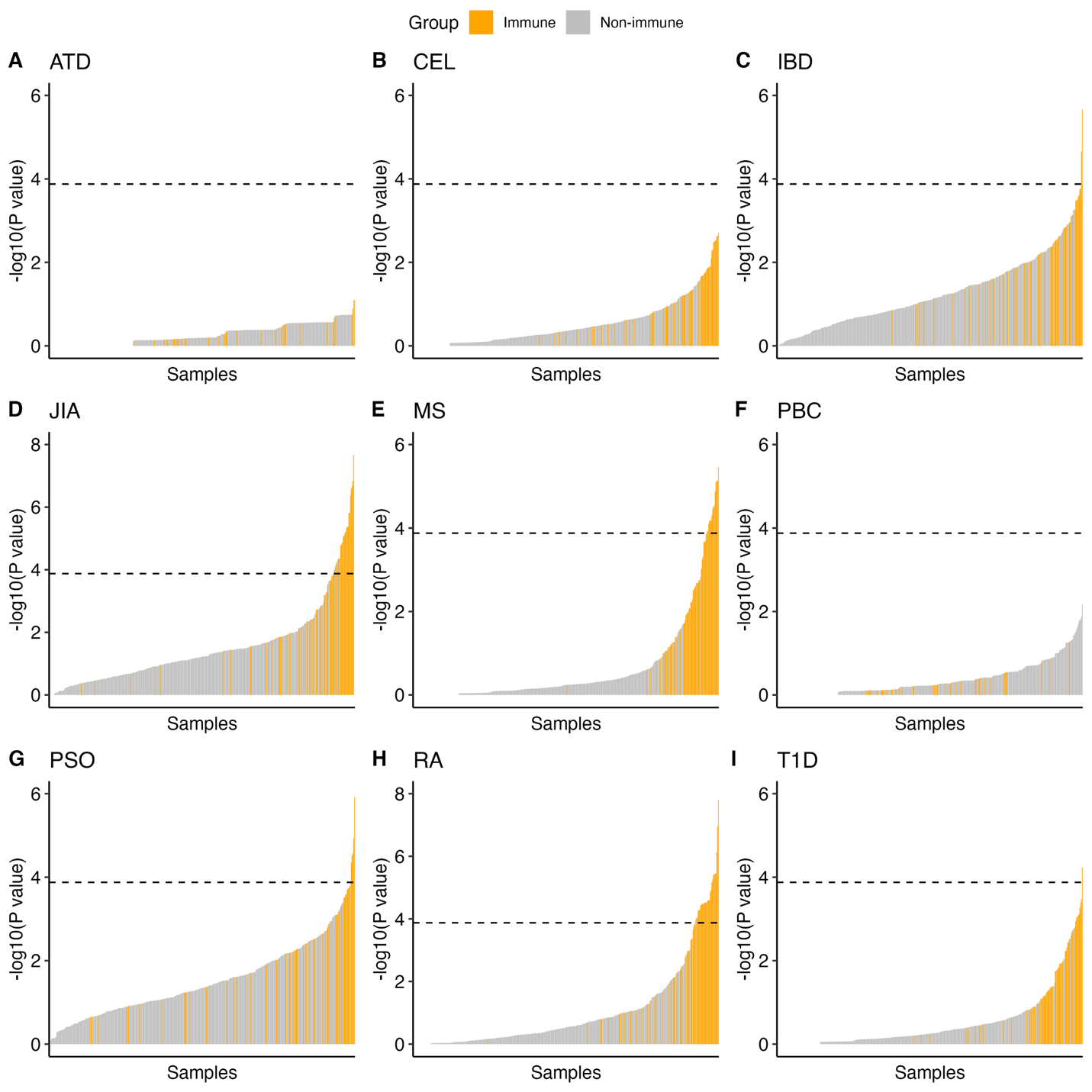

**Supp. Figure S1**: (A-I)barplots showing the -log10 of raw (not corrected) P values of enrichment of samples (each bat represent a sample)  in autoimmune diseases: ATD, CEL, IBD, JIA, MS, PBC, PSO, RA, and T1D. Bonferroni threshold of 5 percent is shown as dashed lines in figures.

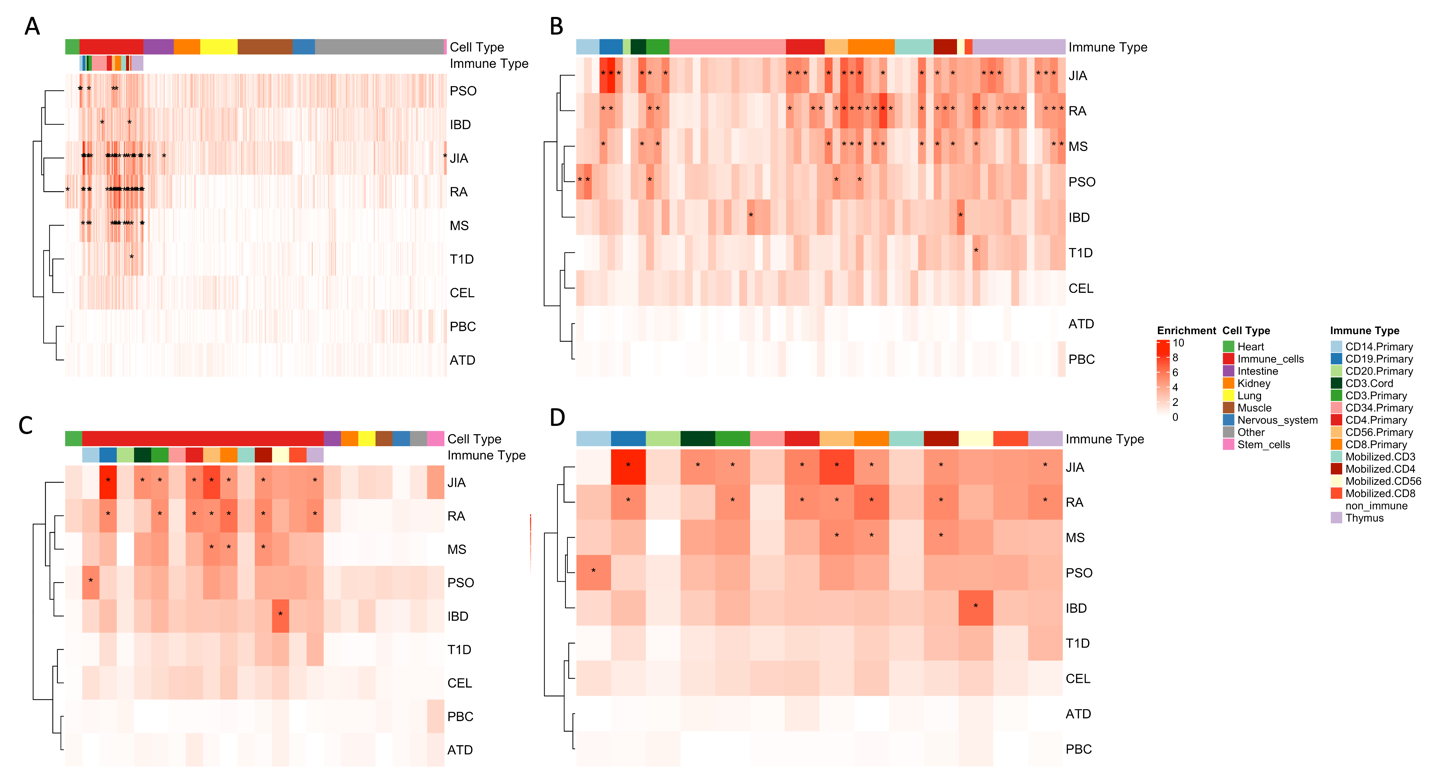

**Supp. Figure S2**: YOU NEED A TITLE HERE. (A) heatmap showing the -log 10 of raw P values of enrichment of samples for each trait. Cell types are specified with different colors in the top colorbar and each row represents one autoimmune disease. (B) heatmap similar to (A) but with only immune cell types. (C) heatmap showing the median enrichment value (-log10 of P value) of samples from a cell type for all cell types and autoimmune diseases. (D) heatmap similar to (C) but with only immune cell types. In all four heatmaps, significant elements (passing bonferroni threshold of 5 percent) are marked with stars.

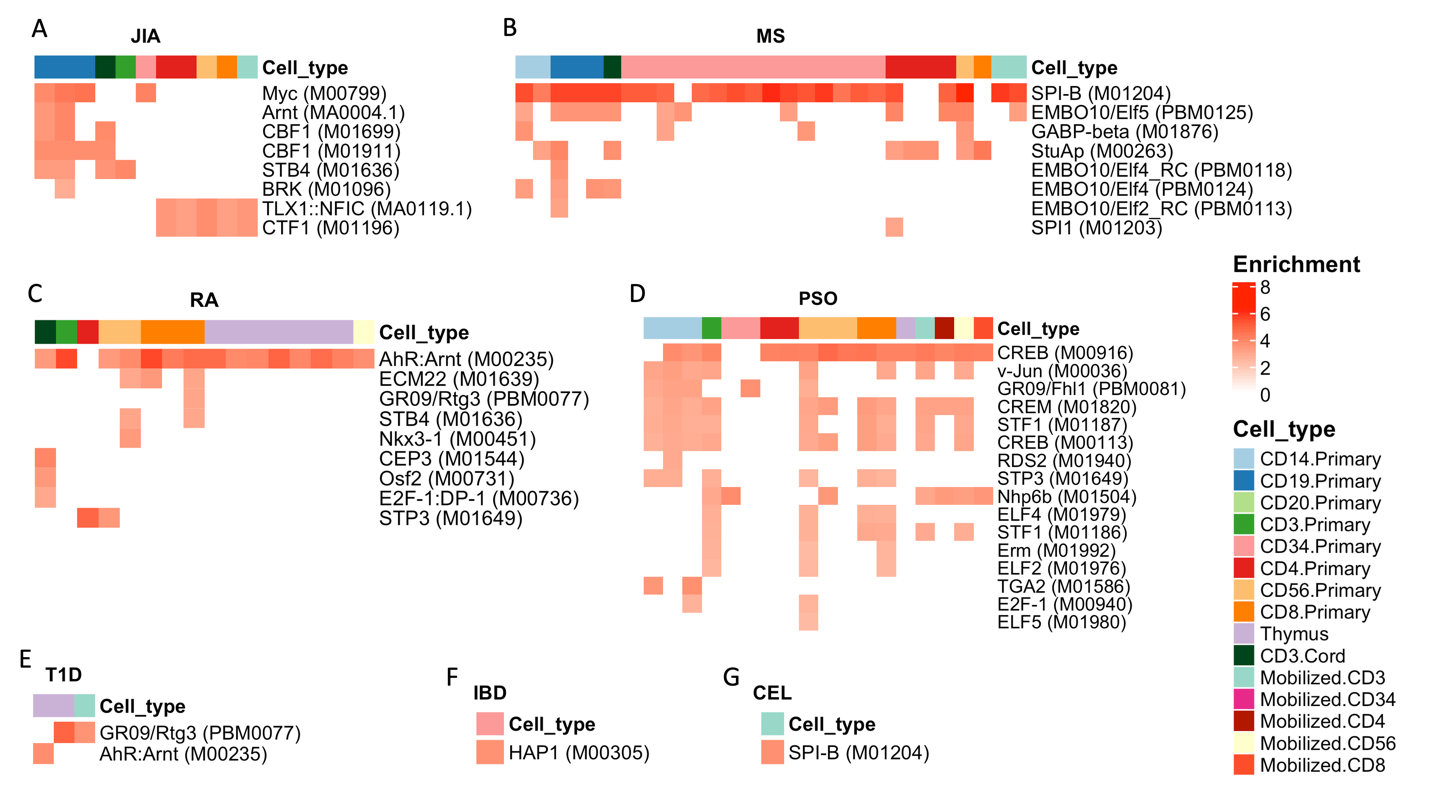

**Supp. Figure S3**: (A-G) heatmaps showing  -log10 of the original P values of enrichment of significant transcription factors (at FDR of 5 percent)  in immune-related samples and cell types for seven autoimmune diseases: JIA, MS, RA, PSO, T1D, IBD, and CEL. In each heatmap, the top colorbar represents the immune cell types.

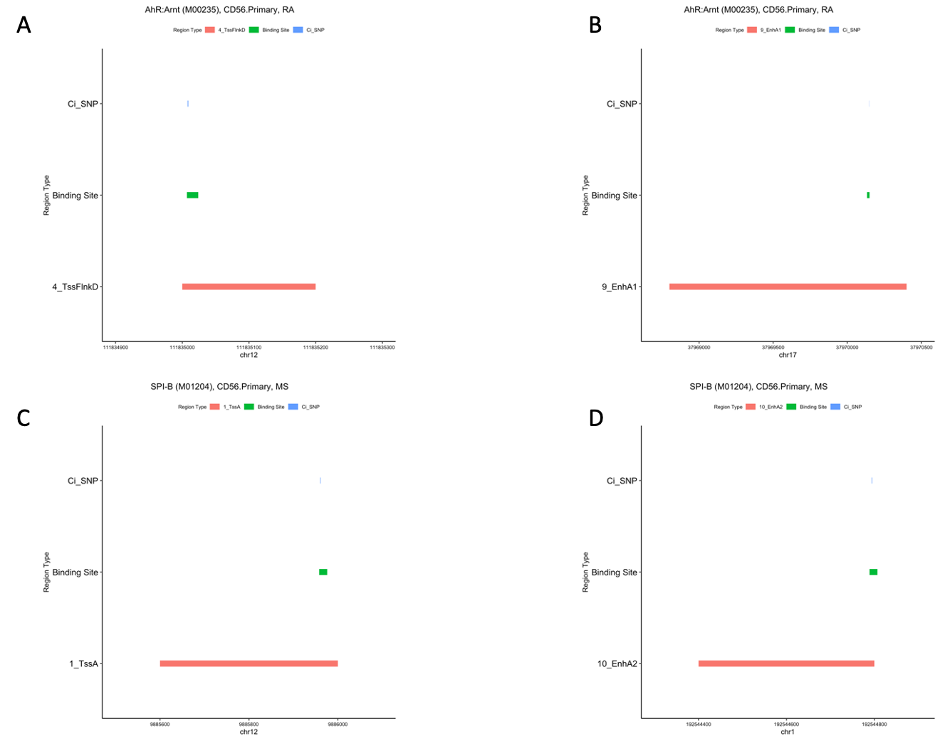

**Supp. Figure S4**: (A,B) segment plots showing binding sites of Ahr:Arnt (most relevant transcription factor to RA), CI SNPs of RA colocalizing to its binding sites, Transcription Start Site Flanking (A) and Enhancer regions (B) overlapping its binding sites in CD56 cell type. (C,D) segment plots of binding sites of SPI-B (most relevant transcription factor to MS), CI SNPs of MS colocalizing to its binding sites, and Transcription Start Site (C) and Enhancer regions (D) overlapping its binding sites in CD56 cell type.

**Titles of Supplementary Tables**

**Supp. Table S1:** Contingency table of the enrichment of risk variants of autoimmune diseases in each sample.

**Supp. Table S2:** Contingency table of the enrichment of risk variants of autoimmune diseases in each transcription factor.

**Supp. Table S3:** Reference cell types from the Roadmap Epigenome dataset corresponding to immune cell types in our data, the cell types that are present in our dataset are marked as used.

**Supp. Table S4:** Table showing the chromosome, start, end of the affected transcription factors, id of the motifs, transcription factor names, frequency of the transcription factors in samples and unique cell types, chromosome, start, end of the effect SNPs affecting them, samples and cell types including them, P values of enrichment of transcription factor and sample, and the autoimmune diseaseAID of interest.

**Supp. Table S5:** The table showing the genes involved in the autoimmune disease-related biological pathways with FDR < 1% and found to be related to autoimmune disease-related transcription factors. This table comprises three columns showing the biological pathways related to autoimmune diseases, their binomial adjusted P values, the genes involved in those pathways, and the autoimmune disease of interest.

**Supp. Table S6:** Transcription factor-gene pairs affected by autoimmune diseases’ risk variants and their cell type activities. This table shows the chromosome, start, end of the binding sites of the affected transcription factors, the cell types comprising those transcription factors, the genes under their control, and their distance to the transcription start site (TSS) of their affected genes.

**Supp. Table S7:** Summary of the data used in the studies that provide risk variants of autoimmune diseases, including number of SNPs, number of cases and controls, and the reference to each study.
